# Phospholipids that plug the pores of cholesteryl ester transfer protein control its lipid transfer dynamics

**DOI:** 10.1101/2025.09.04.674379

**Authors:** Sukriti Sacher, Praveen Singh, Abhishek Mukherjee, Akash Kumar Bhaskar, Kausik Chakraborty, Laurent Counillon, Shantanu Sengupta, Mallorie Poet, Arjun Ray

## Abstract

The mechanism by which cholesteryl ester transfer protein (CETP), a key drug target in cardiovascular disease, moves lipids between lipoproteins, has remained elusive. CETP is the primary protein that regulates plasma circulating levels of lipids (within high, low, and very low-density lipoproteins. The protein, in addition to its terminal openings, has two additional openings, plugged by two phospholipids (PLs). Small molecule inhibitors targeting CETP tunnel displace PL during CETP inhibition. Here, using steered molecular dynamics simulations followed by in-vitro mutagenesis we show that CETP-bound PLs are indispensable in establishing the optimal architecture of CETP tunnel. Our structural and functional analyses revealed that lipid traversal through CETP’s central tunnel is facilitated through hydrophobic interaction-mediated diffusion. PLs are critical in synchronizing domain movements of CETP while accelerating triglyceride traversal through the tunnel and their activity regulated through salt-bridge interactions. Most notably, we show for the first time that phospholipids (PLs) bound within CETP’s tunnel accelerate lipid movement through a novel “gliding” mechanism, representing the first known example of a lipid regulating the movement of another. We identified conserved phenylalanine (Phe) flaps that act as a regulatory valve, opening and closing concertedly to prevent lipid backflow in the absence of an active motor. This study provides in-depth understanding of the mechanism of lipid exchange by CETP guided, and accentuated by its interaction with PLs. The conservation of these critical structural elements across the BPI/LBP family suggests that this mechanism is broadly applicable, expanding our understanding of lipid transport in this clinically significant protein class.

**Significance Statement:** The mechanism of lipid transport by the cholesteryl ester transfer protein (CETP), a major drug target, has long been a scientific mystery. Neutral lipid exchange through CETP is dependent on the hydrophobic characteristics of the tunnel that has been exploited during inhibitor design. We provide a detailed, molecular-level explanation, showing for the first time that phospholipids (PLs), previously thought to be static plugs, are indispensable for lipid transfer. They actively accelerate lipid movement through a novel gliding mechanism, the first documented instance of a lipid regulating another. This discovery identifies new therapeutic targets for CETP inhibition. Our findings also suggest that this fundamental mechanism is conserved across a large protein family, with broad implications for understanding lipid transport and function.

## Introduction

Cholesteryl ester transfer protein (CETP) is a plasma glycoprotein uniquely responsible for bidirectional transfer of cholesteryl esters (CEs) from high-density lipoproteins (HDLs) in exchange for triglycerides (TGs) from low-, intermediate-, and very-low-density lipoproteins (1). This reciprocal exchange depletes HDL of CEs while enriching it with TGs, accelerating its clearance from circulation (2-4). CETP is an active target for therapeutic intervention in cardiovascular disease, owing to its pro-atherogenic role in modulating plasma lipoprotein composition (5-7). However, clinical development has been fraught with setbacks: torcetrapib (a CETP inhibitor) administration to CVD patients increased plasma HDL levels by 72.1% with a 24.9 % decrease in plasma LDL levels (8). However, a significant increase in mortality occurred following its treatment, which was eventually attributed to off-target toxicity. The subsequent inhibitors dalcetrapib and evacetrapib showed insufficient effectiveness (5,9). Later, anacetrapib showed therapeutic effectiveness by lowering major coronary events; nevertheless, it was pulled from development due to its prolonged half-life (5,10). These failures underscore the critical need for elucidating the mechanistic underpinnings of CETP to improve therapeutic interventions.

Two mechanistic paradigms have been proposed for CETP-mediated neutral lipid transfer: a shuttle model, wherein CETP transiently associates with individual lipoproteins (4) and a bridge or tunnel model, in which CETP simultaneously interacts with two lipoproteins, enabling direct lipid flow through its elongated hydrophobic tunnel (4). Electron microscopy supports the formation of both binary and ternary CETP–lipoprotein complexes (11,12); hence, resolving the precise molecular mechanism at high resolution is challenging. CETP contains a continuous hydrophobic tunnel, made by N- and C-terminal β-barrel domains, both of which associate with lipoproteins (11-14). The N-terminal barrel preferentially engages HDL, whereas the C-terminal barrel interacts with TG-rich lipoproteins (11,12). Lipid entry occurs via the open termini and is governed by concentration gradients across lipoprotein surfaces (15). The CETP tunnel can accommodate two CE (16), or two TG molecules (17). The tunnel is occluded in the middle by two phospholipid (PL) molecules, 1,2-dioleoyl-sn-glycero-3-phosphocholine (DOPC), embedded such that their hydrophobic tails are sequestered within the tunnel, and their polar headgroups remain solvent-exposed (PDB ID: 2OBD and 4EWS) (14,16). Earlier these PLs were thought to be exchanged during lipid transport (16), however, experiments with CETP transgene into *PLTP* knockout mice revealed a lack of CETP PL transfer activity (18) proposing PLs plugging CETP as important structural elements that shield tunnel from aqueous exposure (18, 19). However, PL entry into the CETP tunnel and its structural rearrangement (particularly its orientation) in its binding pocket remain inexplicable. Additionally, bacterial permeability-increasing protein (BPI), a structurally homologous protein of CETP, also harbors PLs within its concave groove with a similar orientation as CETP (20), implying a conserved architectural role. MD simulation studies of CETP bound to PLs has suggested that these PLs may influence the conformational dynamics of CETP (21,22); however, their functional relevance to lipid transfer remains unresolved.

CETP activity has been well studied using in-vitro biochemical assays that quantify mass CE or TG transfer from donor HDL/LDL/liposomes to acceptor particles (15, 23, 24). As these assays study end-point kinetic measurements, they quantify the rate of CE/TG transfer as a bulk attribute rather than providing atomistic insights into the neutral lipid transfer process. Furthermore, electron microscope imaging has described CETP binding to lipoproteins at a single-particle resolution and aided in the monitoring of changes in lipoprotein size due to CETP-mediated remodeling (11, 12, 25, 26). However, the resolution of these methods makes them unsuitable for dynamic characterization of lipid transport through CETP. Due to the limitations of standard biophysical techniques, molecular dynamics (MD) simulations are the preferred tool for evaluating lipid transport via CETP. Classical and coarse-grained MD simulations of CETP-HDL (binary) complex have highlighted the role of terminal W105, W106 and W162 in anchoring CETP to HDL, while F35, F93, F147 in facilitating CE entry into CETP tunnel (19). Subsequently, steered MD has demonstrated complete CE translocation through the CETP tunnel by modelling a minimal ternary complex comprising of CETP embedded within lipid monolayers (27). The study identified important physical contacts between F115, R158 and F167 in the N-barrel domain of CETP with CE allowing it to enter the tunnel; I15, L23, A202, I205, L206, F263, F265 and M433 in the neck region of CETP that act as an energy barrier during CE transit; and F301, M412 in the C-barrel region of CETP that allow for the eventual exit of CE from the tunnel. The CETP system modelled in this study excluded bound PLs, overlooking a critical steric and biochemical constraint. While CE movement through CETP is well characterized (19, 22, 27), TG movement through CETP tunnel has not been studied to date.

The mechanism of CETP inhibition by small molecules (torcetrapib, anacetrapib) has been elucidated by steering the movement of CE and calculating the free energy changes in the presence of inhibitor (28). These observations have suggested that CETP inhibitors physically occlude the tunnel leading to CE taking a different path to move out of the tunnel under a steering force (28). Interestingly, CETP inhibitors displace PL during binding (25), however, the effect this has on CETP lipid dynamics remains unresolved.

To address these deficiencies, we performed extensive classical MD and SMD simulations of CETP, bound and unbound, to PLs, to elucidate the mechanism of TG transport through CETP (**Fig. 1*A***) followed by in-vitro site directed mutagenesis. Our simulations established a pivotal structural role for the embedded PLs in stabilizing the tunnel’s hydrophobic milieu, preventing the loss of lipid substrate and facilitating efficient lipid passage, through powerstroke-like accelerating movements. Further, these PLs promoted synchrony between N- and C-terminal domains facilitating ternary complex formation. We have also identified several critical residues that guide the movement of TG and regulate PL-mediated CETP activity. In turn this advanced our fundamental understanding of lipid transfer within the BPI/LBP (bacterial permeability increasing/lipopolysaccharide binding protein family and provided a mechanistic foundation for designing next-generation CETP-targeted therapeutics.

**Fig. 1.**
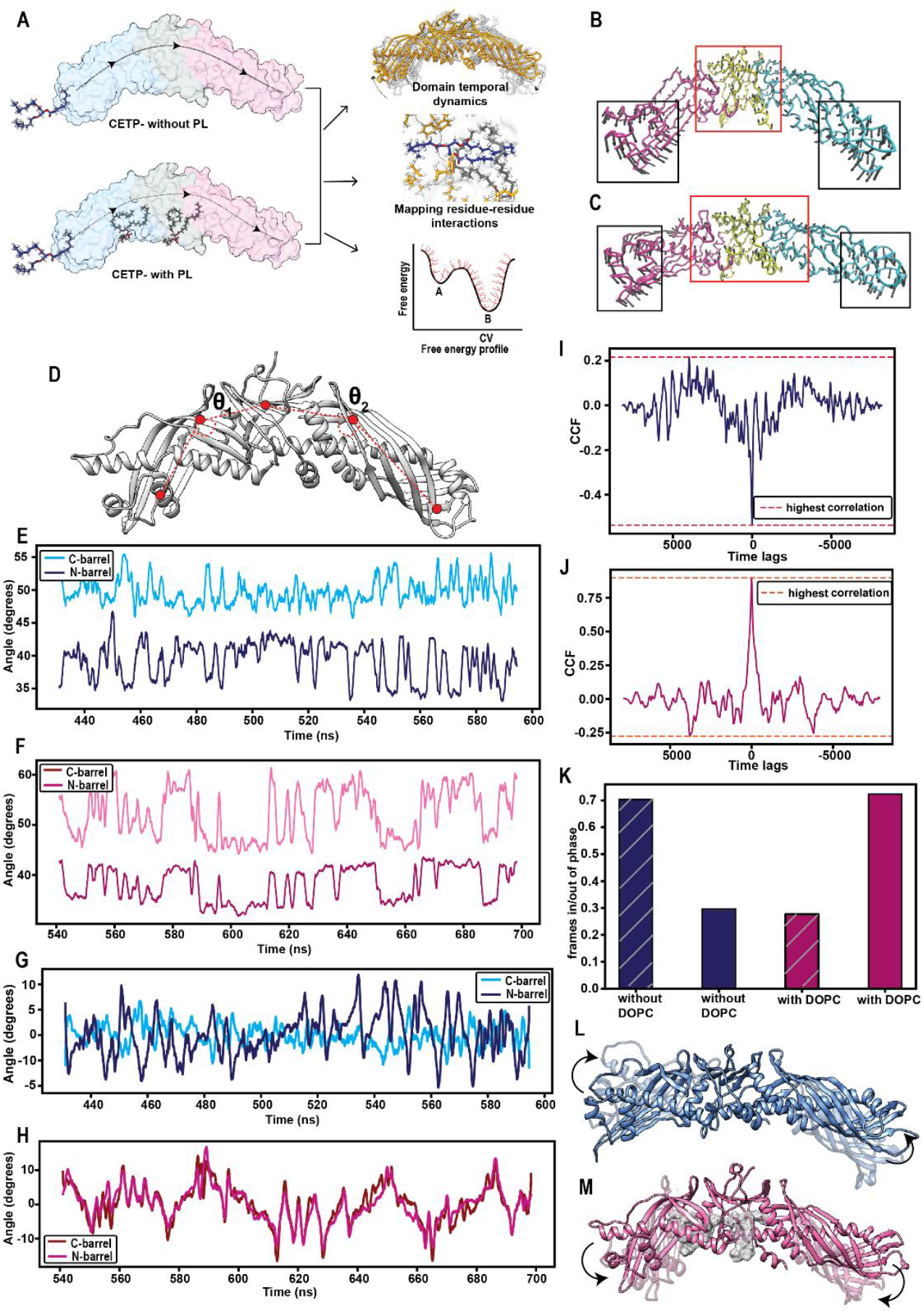
PL plugs results in a switch in synchronicity of oscillations of CETP barrels. **(A)** Schematic of the study design. **(B)** Normal model analysis depicting the two dominant modes of motion in absence (top) and presence of PL plugs (bottom). The length of the arrows depicts amplitude and the arrowheads depict the direction of motion. The protein is colored according to its three domains: C-barrel (pink), Neck (yellow) and N-barrel (blue). **(C)** vectors used to compute barrel movement. Dynamic movements of the C-barrel and N-barrel domains of CETP in the **(D)** absence and **(E)** presence of PL plugs. Rate of change of phase angle of the time series representing the movement of C-barrel and N-barrel in **(F)** absence and **(G)** presence of PL plugs. Cross-correlation between movements of the C-barrel and N-barrel in the **(H)** absence and (**I)** presence of PL plugs. **(J)** Duration of simulation, in which the two barrels remain synchronous (in-phase) or asynchronous (out-phase) in the presence or absence of PL plugs, respectively. The duration is depicted as a ratio of the number of frames in/out phase across triplicate simulations to the total number of frames. Schematic representation of **(K)** asynchronous and **(L)** synchronous movement of the CETP barrels.

## Results

The structure of CETP (PDB ID: 2OBD) was simulated with and without its DOPC plugs for deciphering the role of PL plugs in lipid transfer. During unrestrained simulation, the root mean square deviations plateaued 3–5 and 2–3 Å for CETP simulations with or without PL, respectively. Structures sampled post-stabilization showed a significant increment in tunnel volume within the neck and adjacent N-barrel region of CETP in presence of PLs while tunnel hydrophobicity remained unchanged in both systems.

### PL plugs synchronize the movement of CETP barrels

In the crystal structure, the two barrels of CETP appear bent, providing it a boomerang shape (16). However, the average tilt of the CETP barrels post simulation differed drastically (47.5° and 45° in the absence and presence of PLs, respectively) from that of the crystal structure (72.9°). Normal mode analysis indicated that the terminal regions of the barrel exhibited large movements towards each other irrespective of PL-plug status of CETP (**Fig. 1B and C)**. However, in the presence of PLs, the neck and its adjacent regions, in addition to the barrel ends, exhibited small movements (**Fig. 1C**).

The differential motion of CETP in the presence and absence of PLs intrigued us to investigate the rhythmicity within barrel motions. We calculated the angle formed by the N-barrel and C-barrel with respect to the neck across simulation time (**Fig. 1D**). The N-barrel oscillated between 35–40°, irrespective of the PL status (**Fig. 1E and F**). In contrast, the angle of C-barrel oscillations showed mean fluctuations of approximately 10° (between 47–57°) in the presence of PLs compared to 2° (between 50–52°) in the absence of PLs (**Fig. 1E and F**). Further, the movements of N-barrel and C-barrel negatively correlated in the absence of PLs (**Fig. 1I**), whereas they positively correlated when the PL plugs were present (cross correlation coefficient, 0.8 at 0 lags) (**Fig. 1J**). This implies that the movements of N-barrel and C-barrel in CETP with PLs are temporally similar, and movement of the two barrels are synchronized with each other. Movements of the barrels about a mean position can be considered a periodic function [x(t)] as they return to their respective equilibrium positions at regular intervals; however, the speed of barrel displacement (represented by frequency in a periodic signal) and exact angle of displacement of the barrels (represented by the phase of periodic signal) may vary. To determine these quantities, we performed a Hilbert transformation [x^**^**^(t)] of the angle–time series [x(t)]. On overlaying x^**^**^(t) of the two barrels, the rates of their phase change varied in the absence of PLs (**Fig. 1G**). In contrast, the two barrels exhibited synchronous changes in phase in the presence of PL plugs (**Fig.1H**). The number of frames exhibiting such concerted movement was much higher (> 70% of simulation time) in the presence of PLs than in their absence (**Fig. 1K**) implying that the movement of the two barrels was synchronized in the presence of PL plugs (**Fig. 1M**) than in their absence (**Fig. 1L**).

### PL plugs alter the route taken by TG ensuring successful traversal through the tunnel

To study the TG transfer dynamics through CETP tunnel, we placed a TG molecule at the mouth of the C-barrel and performed SMD on these two systems in the absence or presence of PLs (5,6). A pull was considered successful if TG entered from the C-barrel and exited through the N-barrel end after traversing the entire tunnel. Next, we identified all the hydrophobic clusters lining the tunnel that TG encounters during transit (**Fig. *2A* and *B***). The paths taken by TG clustered together in the presence of PLs, whereas they were scattered (specifically in the two-barrel regions) when PLs were absent (**Fig. 2A and B**). A comparison between the average path taken by TG and the previously identified path for CE (27, 28) indicated that it interacted with 18 common residues (I15, L23, F35, F93, V198, I205, L206, F263, F265, F270, Y361, F363, M412, F408, F429, L425, F429, and M433.

**Fig. 2.**
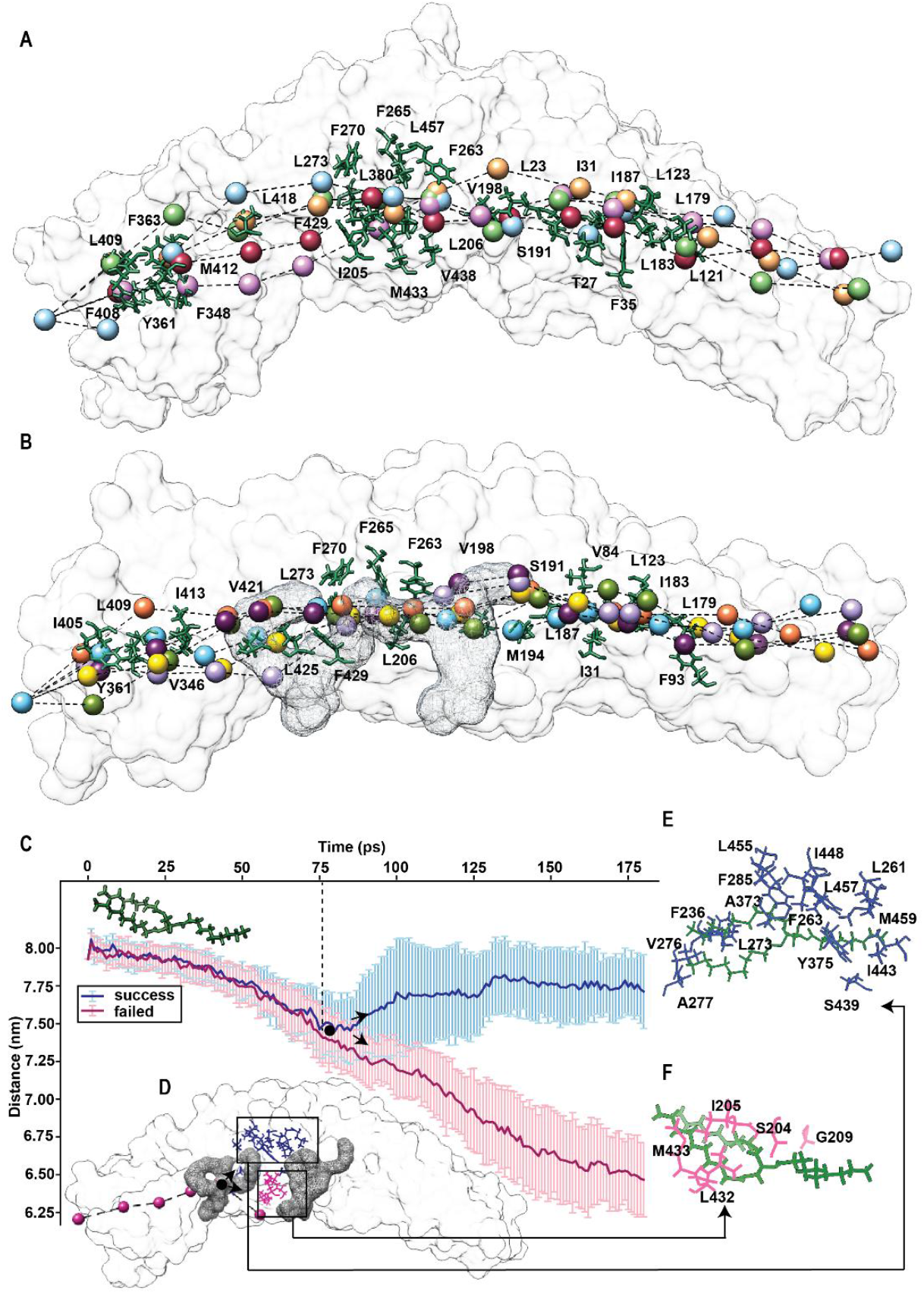
TG transfer through CETP. The center of mass of the glycerol moiety of TG was traced as it moved through the tunnel, in each replicate simulation of CETP **(A)** without and **(B)** with PLs determining the path taken by TG through the CETP tunnel. The residues within a radius of 0.3– 0.6 nm that stay in contact TG for > 1 ns of simulation time in six replicates is highlighted. Multiple paths derived from several independent replicate simulations are depicted. (**C**) The average path taken by TG (across 43 replicate simulations) either directs it through the neck (blue) or leads it out into the milieu through the space in the floor of CETP (pink). Error bars depict standard deviations. The inflection point is depicted as a black circle. The inset shows the path taken by TG, which leads to its fall out, and location of PL plugs on CETP.

In the absence of PLs, although TG entered the tunnel via the C-barrel end (seven instances), it exited downwards through the space between the C-barrel and neck in two of the replicates. We revisited these failed simulations for elucidating the cause of this premature exit. Starting with a simulation frame such that TG had reached the end of C-barrel, TG was pulled toward the N-barrel with random velocities. In 51% of these simulations (43 replicates), TG dropped out through the space between the barrel and neck when PL plugs were absent (**Fig. 2C**). However, not a single fall was observed in the presence of PLs. Next, we calculated the residues that made a contact with TG and identified two hydrophobic patches towards which TG gravitated. The first patch was located towards the roof of the neck region and comprised residues F236, L261, F263, F265, L273, V276, A277, A373, Y375, S439, I443, I448, L457, M459, and L485 (**Fig. 2E**). The second patch was located towards the floor of the neck region and comprised residues S204, I205, G209, L432, and M433 (**Fig. 2F**). Therefore, TG navigated through the tunnel by moving from one hydrophobic cluster to another. Since TG encounters hydrophobic interactive forces only, its contacts with tunnel-lining residues are transient and, therefore, easy to break and form, as TG shifts its position. A bifurcating inflection point lies at the interface of the C-barrel and neck, beyond which two fates for TG exist: TG either reaches the center of the neck or falls out through the opening in the floor of the protein (**Fig. 2D**). Interestingly, the first PL plug was located at this inflection point, suggesting that the PLs physically seal CETP such that TG does not slip out during transit. Moreover, favorable interactions with the acyl tails reroute TG such that it always moves towards the hydrophobic patch at the roof of CETP, reducing the probability of fallouts.

### Side-chain motion of the residues lining the neck region allows the passage of TG into the tunnel

Of the residues that made a contact with TG for an extended duration, three Phe residues were uniquely oriented and also extensively interacted with PL acyl tails. These residues (F270, F265, and F263) were located on the roof of CETP and had their side chains buried inside the cavity. Interestingly, these Phe residues are conserved in other members of the BPI family; F265 is found in PLTP; F263 is found in lipopolysaccharide (LPS)-binding protein; and F270 is present in PLTP, LPS-binding protein, and BPI in addition to CETP (**Fig. 3A**).

**Fig. 3.**
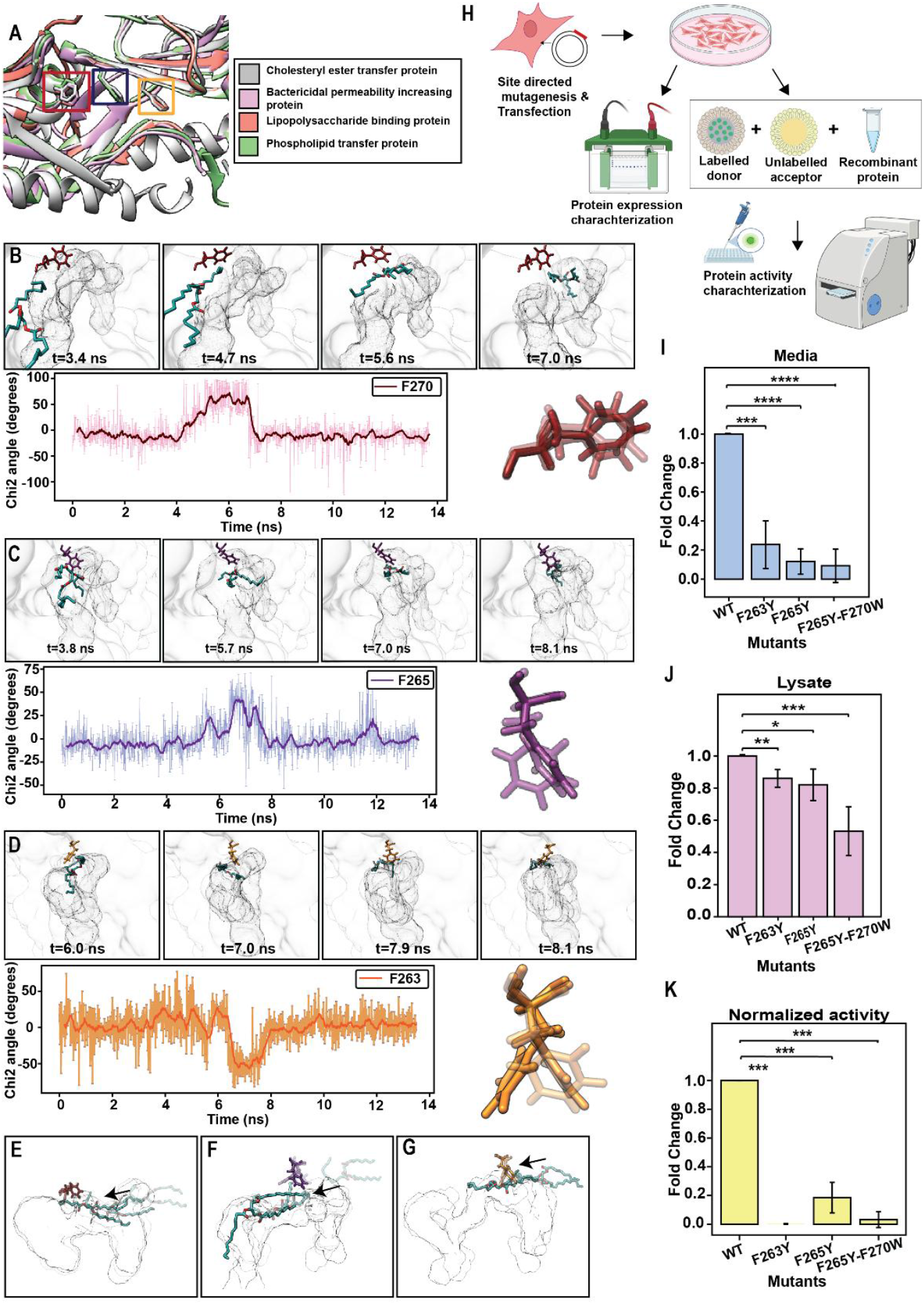
Movement of the flap residues lining the neck region. **(A)** Location of flaps in other members of the BPI/LBP family. PDB structures of CETP (PDB ID: 2OBD), BPI (PDBID: 1EWF), lipopolysaccharide-binding protein (AF-P18428-F1) and PLTP (AF-P55058-F1) overlaid using Chimera. F270, F265, and F263 are highlighted in red, blue, and yellow, respectively. Changes in normalized χ2 angle of **(B)** Phe270, **(C)** Phe265, and **(D)** Phe263 throughout simulation. Snapshots of the simulation depict the movement of Phe side chains as TG approaches it. The solid line depicts the moving average across 20 ps in a simulation. The line plot represents the changes in χ2 angle in real time. Snapshots depicting the movement of **(E)** F270 **(F)** F265 **(G)** F263 when TG was pulled from N-barrel to C-barrel. **(H)** Schematic depicting the study design for experimental validation. Bar plot depicting fold changes in CETP expression of the mutants with respect to that of the WT **(I)** media and **(J)** cell lysate. **(K)** Measurement of CETP activity normalized with respect to protein expression. Values from three biological replicates are shown. The data are presented as mean ± standard deviation. \**p* < 0.05, \*\**p* < 0.01, \*\*\**p* < 0.001, \*\*\*\**p* < 0.0001; ns, not significant (*t*-test).

To ascertain if these residues altered the route of TG, we calculated the torsion angles of their χ^2^ side chains throughout the pulling simulations. These residues underwent frequent rotamer switching irrespective of the PL status of CETP. However, in the presence of PL plugs, as TG slid past these residues, they rotated by approximately 50° to pave the way for TG movement (**Fig. 3B–D**). Once TG passed, the side chains went back to their original orientation. This motion of the three Phe side chains was coordinated, moving sequentially as TG approached them. To test the direction of movement of Phe side chains, we pulled TG in the reverse direction (from the N-barrel to the C-barrel) and observed that the Phe side chains swung open again, allowing TG to pass through (**Fig. 3E-G**).

Given the importance of the Phe flaps, further underscored by their evolutionary conservation, we mutated them to Tyr (F263Y, F265Y, and F265Y-F270Y). HEK293 cells were transfected with plasmids expressing wild type (WT) CETP or mutants, and CETP expression in the media and cell lysate were analyzed using western blotting (**Fig. 3H**). All the three Phe mutants showed poor CETP expression in the media, with F263Y, F265Y, and F265Y-F270Y expression being 0.25-, 0.20-, and 0.1-fold, respectively compared to that of the WT (**Fig. 3I**). For F263Y and F265Y, CETP expression in cell lysate declined by 20%, while the F265Y-F270Y double mutant showed a 50% decline in CETP expression in cell lysate, compared to that of the WT (**Fig. 3J**). These results clearly indicated that the flap (F263Y, F265Y, and F265Y-F270Y) mutants altered CETP structure, affecting either its folding or secretion, thereby leading to decreased expression. Furthermore, the flap mutants presented highly negative effects on CETP activity, with F263Y showing negligible activity, and F265Y and F265Y-F270Y showing > 70% decline in activities (**Fig. 3K**).

### PL plugs facilitate TG movement forward through a power-stroke-like motion

As TG moved through the neck region, past the PL tails, it exhibited a forward thrusting movement. This movement was only apparent in the neck region and not during TG entry or exit through the barrels (**Fig. 4A**). To quantify this thrust, we determined the instantaneous velocity of TG which, increased as TG moved past PL acyl tails (**Fig. 4B**). The occurrence of spikes in the instantaneous velocity coincides with an “assisted gliding” motion. Closer observation reveals the formation of a spatial hydrophobic planar surface due to the interaction between PL tails and TG, allowing it to glide forward (**Fig. 4C**). This gliding is reminiscent of power-stroke motion of actin-myosin, leading to sliding of thin actin filaments over thick myosin filaments (29).

**Fig. 4.**
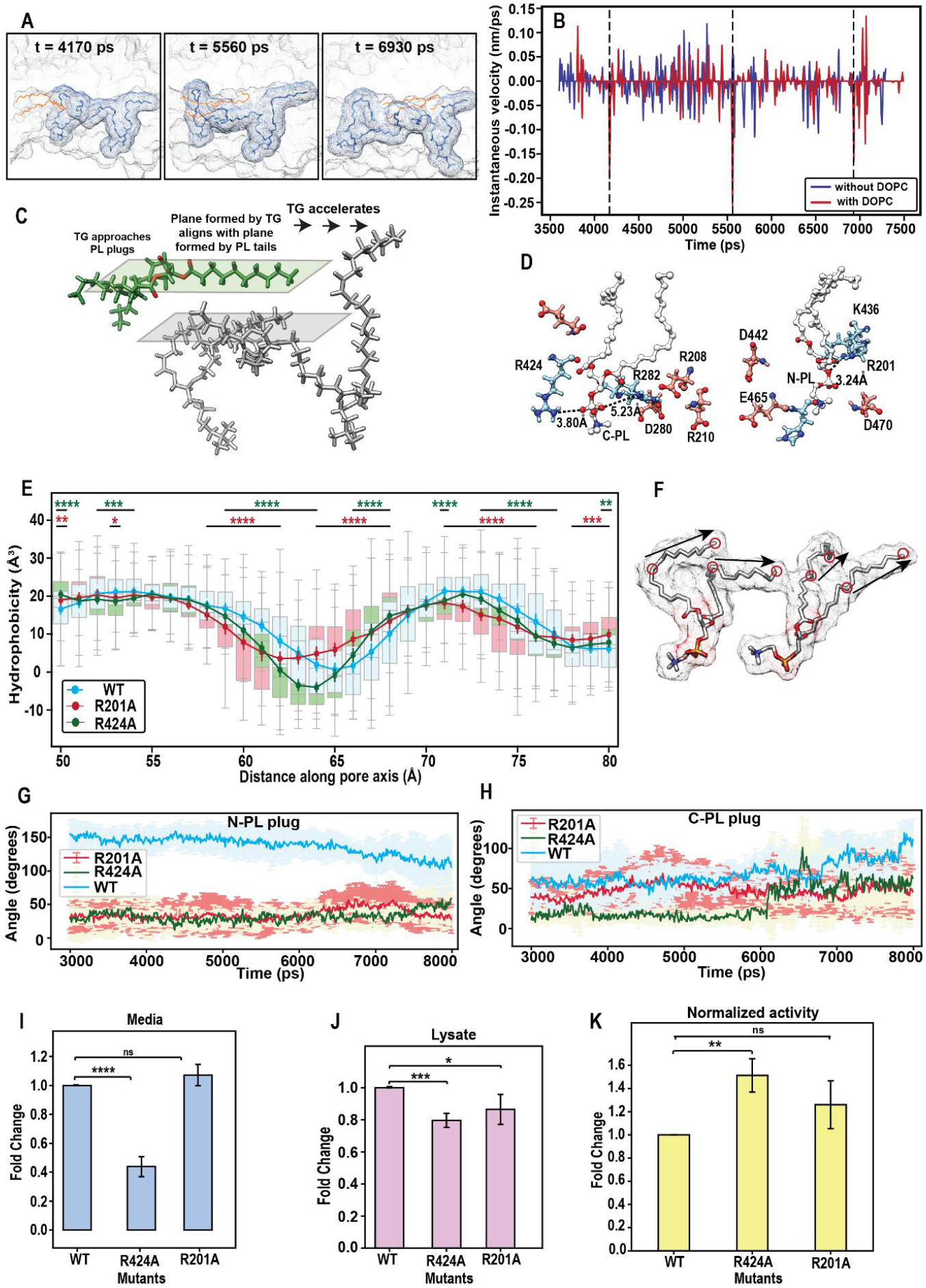
Power-stroke movement of PL acyl tails of PL. **(A)** The time points, at which a forward gliding movement of TG is observed, depicted through simulation snapshots **(B)** Instantaneous velocity of TG through the neck region of CETP **(C)** Schematic depicting the gliding of two hydrophobic planes formed by TG and PL acyl tails (**D**) Salt bridge forming residues around C & N-PL plugs (**E)** Tunnel hydrophobicity of the neck region of WT and mutant CETP. The box plot depicts the distribution of hydrophobicity values calculated over 100 frames (post-stabilization). Solid lines depict the median value of distribution for the distance along the tunnel axis. Mann– Whitney U test was used for statistical significance of the difference between the profiles of WT and R201A (red), and between WT and R424A (green). **(F)** Schematic of vectors defining the movement of PL acyl tails for panel G & H. Angle between the acyl tails of **(G)** N and **(H)** C-PL through SMD simulations. Bar plot depicting fold changes in CETP expression of the mutants with respect to that of the WT in **(I)** media and **(J)** cell lysate **(K)** Measurement of CETP activity normalized with respect to protein expression. Values from three biological replicates are shown. The data are presented as mean ± standard deviation. \**p* < 0.05, \*\**p* < 0.01, \*\*\**p* < 0.001, \*\*\*\**p* < 0.0001; ns, not significant (*t*-test).

To test the validity of the hydrophobic plane’s role is assisting the activity of the protein, we identified the residues in the central pores of CETP that surround PL headgroups. These primarily comprised of charged residues (Arg, Lys, Asp, and Glu) that formed salt bridges with the PL headgroups (**Fig. 4D**). Postulating that the increased rigidity and stability of the plane due to decreased fluctuations of PLs should increase the activity, we mutated R424 and R201 near C-PL (PL between the C-barrel and neck) and N-PL (PL between the N-barrel and neck), respectively to Ala. *In silico* characterization of the two mutants showed increases in tunnel hydrophobicity, specifically in the neck region (**Fig. 4E)** Moreover, the overall fluctuations of PL plugs were reduced in the mutants in comparison to WT CETP. But the extremities of acyl tails exhibited a unique orientation such that they lay parallel to each other (**Fig. 4F-H**) forming a rigid but stable hydrophobic plane (**Fig. 4C**).

These results were further validated by mutating these residues *in vitro*. HEK293 cells transfected with plasmids expressing WT CETP or mutants, were analyzed using western blotting. R201A showed an expression similar to that of the WT in the media. However, R424A expression was significantly reduced (0.4-fold compared to that of the WT in the media) (**Fig. 4I**). CETP expression in cell lysate declined by 20%, for both R201A and R424A, compared to that of the WT CETP (**Fig. 4J**). Furthermore, both R424A and R201A had significantly higher activities than that of WT CETP (**Fig. 4K**). The increased stability of the hydrophobic plane may lead to an increase in power-stroke-like events contributing to fast lipid transfer through the tunnel in these mutants.

### Mechanism of lipid transfer through CETP

Next to deduce the free energy of TG traversal within the CETP tunnel, we calculated the potential of mean force (PMF) by choosing windows (spaced 0.1 nm apart) along the reaction coordinate (RC) defined by the path taken by TG within the tunnel **(Fig. 5A)**. The PMF profile suggested that the entry of TG into CETP tunnel is thermodynamically favorable (with a downhill slope of ∼ - 3.5Kcal/mol-nm) as it moves from an aqueous to a hydrophobic environment. On its entry within the tunnel, it encounters a cluster of mostly hydrophobic residues (L287, L296, E306, I307, V311, F348, F408, M412) which forms a shallow barrier of ∼ +1Kcal/mol. Once past this barrier the movement deeper into C-barrel is stabilizing (gradual descent of -1Kcal/mol). At the interface of C-barrel and neck, a shallow well of depth 1Kcal/mol is observed. The movement across this minimum is guided by interaction with acyl tail of C-PL (RC: -8.1 nm). As TG enters the neck, it undergoes a gradual ascent (slope = 1.2Kcal/mol-nm) in three steps. The rise from the well to the first saddle presents with a small energy barrier of 1Kcal/mol. This represents the opening of F265 gate. The rise to the next saddle represents opening of F263 gate. Finally, the movement past the third saddle into the N-barrel (the narrowest region of the tunnel) is achieved through the gliding hydrophobic interaction between N-PL acyl tail extremities and TG, resulting in a power stroke. It is important to note that the free energy within the neck remains negative, and the local fluctuations of PL tails tug the TG forward inhibiting kinetic trapping within this region. As TG moves further into the N-barrel, closer to exit, it interacts with another cluster of mostly hydrophobic residues (Y40, Y57, L59, I117, V178, W162, F167). Further, the energy barrier increases sharply (steep ascent of 4Kcal/mol) in this region. On comparing it with the deepest point in the PMF (interface of C-barrel & neck), this energy barrier amounts to 5Kcal/mol which is high enough to strongly resist spontaneous passage. The steepness of this barrier is further exacerbated due to the movement of TG from a favorable hydrophobic milieu to aqueous one.

**Fig. 5.**
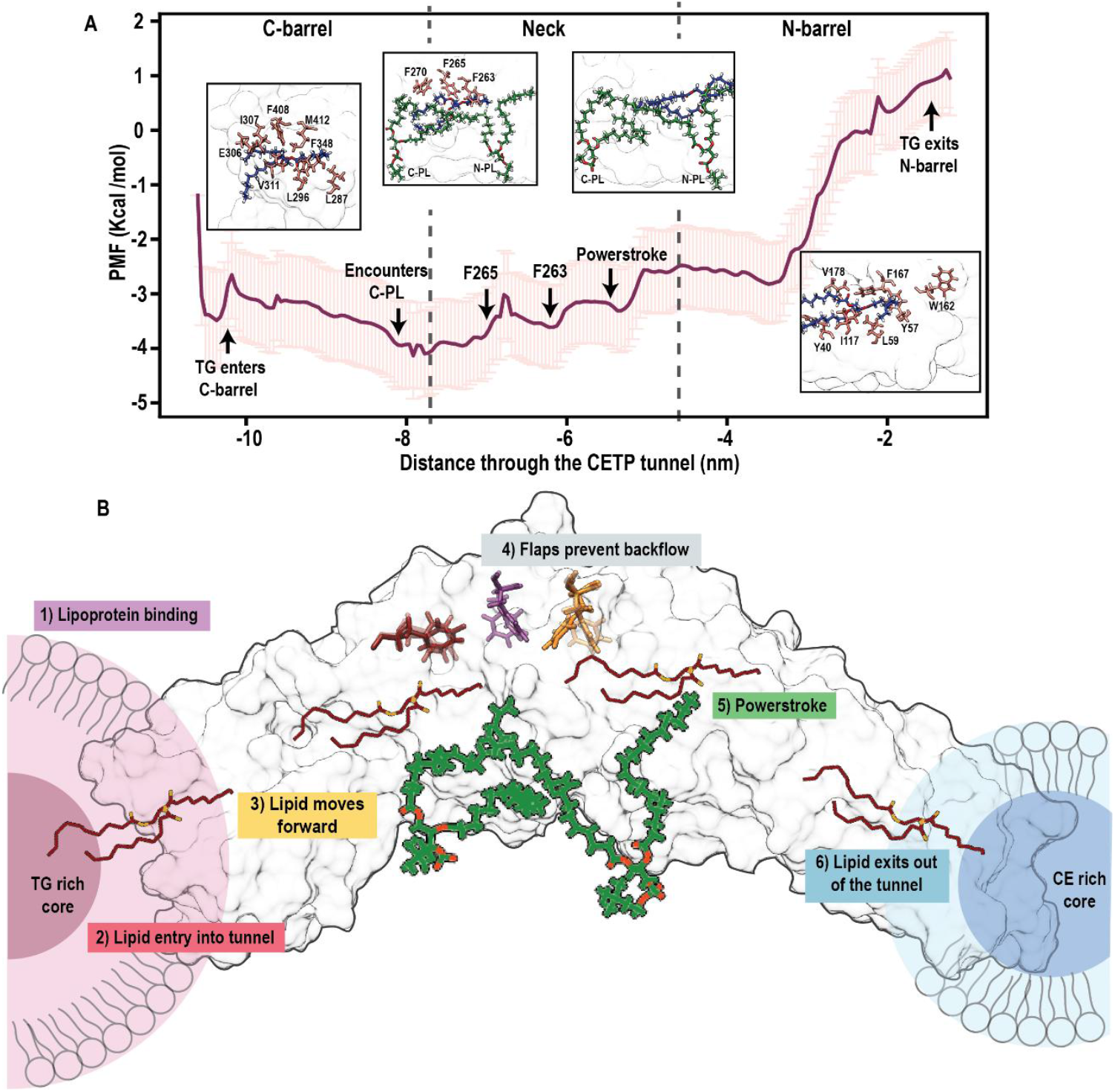
Mechanism of lipid transfer through CETP. **(A)** Potential of mean force profile for the movement of TG through the CETP tunnel. Errors bars depict standard deviations from the mean and are calculated using bootstrapping. The major events that occur in the CETP tunnel are depicted with arrows and snapshots respectively. **(B)** Schematic describing the six steps involved in the mechanism of TG transfer. PL-induced domain synchrony results in binding of CETP to lipoproteins (Step 1). Upon CETP attachment and penetration into lipoprotein surface, neutral lipids present near the surface enter the CETP tunnel owing to a difference in concentration gradients (Step 2). Once inside the tunnel, peristaltic forces generated by domain movements of CETP guide the lipid further into the tunnel. Additionally, transient contacts with the hydrophobic clusters located inside the tunnel steer the lipid forward (Step 3). As an internal check in the neck region, the concerted movement of the flaps ensures no lipid reflux (Step 4). Subsequently, the hydrophobic interactions between PL acyl tails and lipid leads to acceleration of lipid forward (Step 5). On reaching the end of the tunnel, lipid exits owing to a difference in concentration gradients (Step 6).

## Discussion

This study establishes a critical role of PL plugs in modulating the structural and functional dynamics of CETP. PL plugs enhance hydrophobicity of the CETP tunnel by sealing its two central orifices and rerouting TG transport to ensure it is not lost during transit. Furthermore, they promote forward lipid movement through favorable hydrophobic interactions between PL acyl tails and TG. Interestingly, PL acyl tails assist in the conversion of CE from a bent to linear form during its traversal through the neck region of the tunnel (22). This linear CE may similarly glide over a hydrophobic plane formed by PL acyl tails, just as we observed for TG, to produce a forward translational motion. While the role of PLs in shielding lipid cargo from aqueous exposure has been proposed (16,30), their impact on tunnel hydrophobicity and lipid mobility has not been demonstrated prior to this work.

Our study further identifies key structural elements within CETP, which regulate lipid traffic. Three Phe residues (F263, F265, and F270) form flaps at the neck region, which stabilize CETP and modulate lipid passage. Given the location and orientation of these residues in the bottleneck of protein and the distinct closed and open states exhibited by these gates, we believe that they may play a role in controlling the rate of transfer and direction of lipid cargo. Mutation of F265 or F270 to arginine abolishes CETP expression probably owing to misfolding (16), and even substitution with Tyr significantly reduces CETP expression and activity. These residues likely function as directional valves as opposed to hinged gates, allowing both influx and efflux of CE and TG at differing rates (24). Residue gates in proteins ensure substrate selectivity, prevent solvent access to certain regions, and synchronize processes that occur in different protein compartments (31). Similarly, these flaps may prevent regurgitation, such that a substrate, upon entering the tunnel, does not reflux out. Considering the movement of lipid through CETP is entirely driven by diffusion, opening, and closing of these flaps is extremely important to prevent backflow of lipid in the absence of any active motor driven transport. Furthermore, the conserved nature of these residues across BPI/LBP family members may suggest a shared mechanistic role.

Electrostatic interactions around the PL headgroup also affect CETP stability and activity. Missense variants in this region (R282C and D442G) are associated with hyperalphalipoproteinemia characteristic of CETP deficiency (32). Salt bridges forming residues R201 and R424 modulate CETP activity. Interestingly, R201 is also involved in interaction with torcetrapib (CETP inhibitor) and its substitution to alanine, increases CETP activity and IC_50_ of the inhibitor (14). These residues increased favorable hydrophobic interactions with TG; increasing power stroke-like events that push the lipid forward into the tunnel. These residues also increased tunnel hydrophobicity, presenting themselves as additional regulatory points that can influence CETP tunnel and lipid transfer dynamics.

In addition to sealing the tunnel, PLs influence the conformational dynamics of CETP. Their presence induces synchronized motion between the N- and C-terminal β-barrels. These concerted movements, characterized by squeezing (barrels moving toward each other) and stretching (barrels moving apart), may generate peristaltic forces that propel lipids through the tunnel. Such rhythmic dynamics may also facilitate lipoprotein binding. Protein domain motions influence ligand interactions (33,34). Moreover, the ability of CETP to bind a second lipoprotein increases after engaging the first, suggesting allosteric communication between the termini. This effect is markedly reduced by inhibitors that prevent ternary complex formation (25). Notably, inhibitors, such as anacetrapib and torcetrapib, displace the PL located at the N-barrel–neck interface (14). Taken together, the displacement of PLs by inhibitors may potentially disrupt barrel synchrony and impede lipid transfer. These findings underscore the importance of PLs in regulating the mechanical and functional behaviors of CETP.

Given their indispensable role, CETP likely circulates in plasma in a PL-bound form. However, PL incorporation post-synthesis via lipoprotein binding appears unlikely: electron microscopy shows that the PL-binding pocket lies >15 Å from the HDL surface (11,12). Similarly, PL entry via the hydrophobic tunnel followed by their rotation for sealing the orifices is geometrically implausible. Alternatively, a scenario in which CETP acquires PLs through its central concave groove by binding to the lipoprotein (16) does not adequately account for the precise orientation required for tunnel plugging. We therefore hypothesize that PLs are integrated during CETP folding or secretion. Furthermore, PL plugs are also present in bacterial BPI, another BPI/LBP family member (20), suggesting that PL-mediated lipid transport may be a conserved mechanism across the family.

We compared the previously identified path for CE (27,28) with the path of TG and observed both the lipids to travel through the same route. Since, CETP inhibitor (torcetrapib and anacetrapib) occlude the tunnel by interacting with residues C13, H232, F263, A202, and V198, which lay in the path of TG, the binding of inhibitor may affect the movement of TG, just as it affects the movement of CE (13,28). Since both the lipids travel through the same route, and their point and direction of entry or exit has no influence on the path that they take, the difference in the rate of CE and TG transfer previously observed (24) must be due to additional factors such as surface availability of these neutral lipids in lipoproteins (23) or auxiliary proteins that influence binding and stability of CETP with lipoproteins (19). Furthermore, missense single nucleotide variations of several of these TG interacting residues (I11V, V136G, I413V, M422I, R352H, V295E, Y361C, V416G, I64N, I205F, V340G, V311I, V409M, L273P, V323I, V344A, L432I, C13Y, R292C, R352C, L206R, D382H, A373P, V89M, A195D) were predicted as possibly damaging potentially affecting CETP function. These mutations may alter tunnel architecture negatively influencing CETP mediated lipid transfer.

Based on our findings, we propose a multi-step mechanism of lipid transfer **(Fig. 5B)**: (1) PL-induced domain synchrony enables lipoprotein docking and penetration; (2) neutral lipids enter the tunnel via CETP termini, guided by concentration gradients; (3) lipids traverse the tunnel through sequential hydrophobic interactions; The local fluctuations of PL acyl tails gently tug the lipid forward guided by hydrophobic clusters (4) Phe residues at the neck function act as a gating system for preventing backflow; (5) hydrophobic interactions with PL acyl chains generate a power-stroke effect, propelling lipids forward into the N-barrel (the narrowest region of the tunnel); and (6) lipids exit through the distal end, driven by differences in lipid concentrations. Further the PMF showed a very high exit barrier which is strong enough to resist spontaneous diffusive movement of TG. However, the presence of lipoproteins at the ends of CETP may significantly reduce this barrier further supporting the formation of ternary CETP complex during lipid transfer.

Our work provides compelling evidence in favor of the tunnel mechanism of lipid transfer. In summary, we demonstrate that PL plugs act as structural cofactors that enhance the lipid-transport efficiency of CETP by modulating tunnel hydrophobicity and protein conformational dynamics. This PL-mediated mechanism may extend to other BPI/LBP family proteins, such as PLTP, LBP, and BPI, offering a broad framework for understanding lipid-transfer processes. F265, F263, F270 flap residues and salt bridge forming residues are important regulatory points in the CETP structure that influence its stability and activity. While CETP inhibition has focused on tunnel occlusion, requiring hydrophobic drug molecules that can enter the CETP tunnel, design of therapeutics that target the solvent accessible PL-binding pockets may prove to be an exciting alternative.

## Materials and Methods

### Computational methods

#### Molecular Dynamics Simulation setup

MD simulations were performed with the GROMACS (35, 36, 37) v.2022.1 suite. The system consisting of CETP with PLs (PDBID: 2OBD) or CETP without PLs was represented by CHARMM36 (38). All starting structures were subjected to a minimization and equilibration protocol recommended by CHARMM. Three independent trajectories, each of 650 ns at 300 K, were carried out for CETP with PLs (CETP-with DOPC). For CETP without PL (CETP-without DOPC), simulations were carried out for 600 ns at 300 K. Simulations for CETP mutants R201A and R424A were carried out for 700ns at 300K.

#### SMD simulations setup

Following the conventional MD run, the last frame of the trajectory was used as the starting point for SMD simulations. One TG (10:0/10:0/10:0) was manually placed at the opening of the C-barrel of CETP (11). SMD simulations were performed independently on each of the two systems in the absence or presence of PL plugs to elucidate the path of TG transfer. TG was pulled through the CETP tunnel using constant-velocity SMD simulations. The pulling vector was defined on the basis of the center of mass of the steered group, i.e., the tail region of TG and center of mass of CETP tunnel residues in the direction of the N-barrel. Considering the asymmetric shape of CETP, its center of mass was dynamically switched from the C-barrel to the neck and finally N-barrel, depending on the location of the steered group. Therefore, entire pull simulation through the CETP tunnel was executed in four phases. For without and without PL, SMD simulations were performed in seven replicates each. For the CETP mutants (R201A and R424A), SMD simulations were performed in triplicates.

#### Calculation of the PMF

Umbrella sampling simulations were performed on paths taken by TG in the presence of PL plugs, derived from SMD simulations. Starting from the initial position of TG at the mouth of the C-barrel, the path traversed through the tunnel was sampled at every 0.1 nm. A force constant of 500 kJ-mol^-1^-nm^-2^ was used. To ensure sufficient overlap of histograms along the reaction coordinate. Each window was thoroughly equilibrated for 1 ns and subjected to a production run for 50 ns. The histograms obtained were unbiased and combined to produce the PMF using the weighted histogram analysis method (WHAM) by excluding the first 20ns from the trajectory (39). An average PMF profile was generated by considering the mean potential at each point along the reaction coordinate with bootstrapping (100 runs) to generate errors. The first and last bins of the reaction coordinate were assumed to be neighbors for generating a periodic PMF.

### Experimental Methods

#### Plasmids and mutants

WT-CETP plasmid was a kind gift from Thea Bismo Strøm, Department of Medical Genetics, Oslo University Hospital, Norway (40). Using pcDNA3.1-WT-CETP-V5 as a template, five mutants were generated using a QuikChange II Site-Directed Mutagenesis Kit (Agilent Technologies, Santa Clara, CA, USA)) according to the manufacturer’s instructions. Plasmids were isolated using a Plasmid Midi Kit (Macherey-Nagel GmbH & Co. KG, Düren, Germany). Plasmid integrity was verified by Sanger sequencing.

#### Cell culture and transfection

HEK 293-F cells (6 × 10^5^ cells/well) were seeded in a 6-well plate (Nunc, Villebon-sur-Yvette, France) and cultured in Dulbecco’s Modified Eagle Medium (Gibco, Villebon-sur-Yvette, France) supplemented with 10% fetal bovine serum (Sigma Aldrich Chimie, Saint-Quentin-Fallavier, France) and 1% penicillin-streptomycin antibiotic (Gibco, Villebon-sur-Yvette, France). Cells at 50% confluence were transfected with 1 μg DNA using a jetPRIME transfection kit (Sartorius Polyplus SAS, Illkirch-Graffenstaden, France). After 2 h, cells were washed with phosphate-buffered saline (Sigma Aldrich Chimie, Saint-Quentin-Fallavier, France) and were incubated in Freestyle 293 expression media (Gibco, Villebon-sur-Yvette, France). Cells were harvested 48 h post-transfection and were lysed using RIPA lysis buffer (NaCl 150mM, 1% d’IGEPAL^®^ CA-630, 0.5% Sodium deoxycholate, 0.1% Sodium Dodecyl Sulfate, Tris 50mM, pH 8) containing a protease inhibitor cocktail (Protease and phosphate inhibitor, Sigma Aldrich Chimie, Saint-Quentin-Fallavier, France). The spent medium was centrifuged at 1200 rpm for 5 min to remove cell debris. Protein concentrations of the lysates and supernatants were determined using RC DC Protein Assay (Bio-Rad Laboratories, Inc., France).

#### Western blotting

Equal amounts (30 μg) of protein samples were separated by 10% sodium dodecyl sulfate–polyacrylamide gel electrophoresis. Prestained Opti-Protein Marker (Applied Biological Materials Inc., Richmond, BC, Canada) was used as a molecular-weight marker. Separated proteins were blotted on a polyvinyl difluoride membrane (Amersham Hybond, 0.2mm. Sigma Aldrich Chimie, Saint-Quentin-Fallavier, France). The membrane was blocked using skimmed milk for 1 h at 25 ℃. Subsequently, the membrane was incubated with an anti-V5 antibody (Invitrogen, Villebon-sur-Yvette, France, R96025) (1:1500 for supernatant and 1:5000 for lysate) for detecting V5-tagged CETP, or anti-β-actin antibody (Sigma Aldrich Chimie, Saint-Quentin-Fallavier, France A5441) which was used as an internal control for cell lysate. An anti-mouse IgG conjugated with horseradish peroxidase (Goat anti-mouse IgG(H+L) secondary antibody HRP, Thermo Scientific #32430, Villebon-sur-Yvette, France) was used as a secondary antibody. Immunoreactive protein bands were detected using (Immobilon Western Chemiluminescence HRP substrate, Millipore) were captured on Fusion Fx (Vilber Lourmat, Marne-la-Vallée, France). CETP expression was analyzed using ImageJ v.1.54p and normalized with respect to β-actin expression^62^.

#### CETP activity assay

Lipid transfer activity of CETP was measured using 4 μg protein from each supernatant (WT or mutant), using a CETP Activity Assay Kit (MAK106; Sigma Aldrich Chimie, Saint-Quentin-Fallavier, France) according to the manufacturer’s instructions. Briefly, fluorescently-labelled donor particles were incubated with the supernatant and acceptor particles in a sealed black flat-bottomed 96-well plate (View plate 96 black, clear bottom, Revity,) for 3 h at 37 °C in a water bath. Fluorescence intensities were measured using excitation and emission wavelengths of 465 and 535 nm, respectively, on a spectrofluorometer (SAFAS Xenius, Monaco) at 400 V.

## Acknowledgments

We thank Katrine Bjune and Åsa Schawlann Ølnes for providing the WT-CETP-V5 plasmid. We also thank Prof. Anand Srivastava (IISc-Bangalore) and Dr. Csaba Daday (Qsimulate) for their insights that helped in shaping this manuscript.

This study was funded by Technology Innovation Hub-IHUB Anubhuti (IHUB Anubhuti/ProjectGrant/25), Science and Engineering Research Board (SUR/2022/005466), and Department of Biotechnology, Ministry of Science and Technology, Government of India (BT/PR45830/MED/12/963/2022). S.Sa. was supported by the University Grants Commission (191620127165), Indo-French Centre for the Promotion of Advanced Research (IFC//4152/RCF-IN-0052, 2023), and Franco India Campus and Overseas Research Fellowship (IIIT-D/ACAD/-1262).

## Author Contributions

A.R., S.Sg and M.P. designed the research. S.Sa. performed the research and analyzed data. L.C., K.C., P.S., A.K.B. and A.M. contributed new reagents. S.Sa, A.R., A.M, S.Sg, M.P. wrote the manuscript. All authors reviewed the manuscript.

